# Deep Learning-Driven Discovery of FDA-Approved BCL2 Inhibitors: In Silico Analysis Using a Deep Generative Model NeuralPlexer for Drug Repurposing in Cancer Treatment

**DOI:** 10.1101/2024.07.15.603544

**Authors:** Ehsan Sayyah, Hüseyin Tunç, Serdar Durdağı

**Author notes:** (SD); (HT).

## Abstract

Finding strong inhibitors of the BCL2 target, which is essential for controlling apoptosis and ensuring the survival of cancer cells, has prompted research into FDA-approved drugs. This study uses an advanced deep generative model called NeuralPlexer to produce protein-ligand complex conformations one-by-one and perform *in silico* analysis. This is the first time in the literature using NeuralPlexer in virtual screening, as we comprehensively evaluate the conformations of an FDA-approved drug library to ascertain their potential efficacy in suppressing BCL2 by utilizing NeuralPlexer’s capabilities. Obtained results were re-confirmed by physics-based molecular simulations and neural relational inference (NRI) analysis. Our study reveals several intriguing candidates such as Lathyrol and Fadrozole with potent inhibitory interactions with the BCL2 target, offering important new information for repurposing currently available drugs in cancer treatment. This work highlights the promise of deep learning technology in pharmaceutical research by integrating NeuralPlexer into the process of drug development, while also improving the accuracy of predictions made about protein-ligand interactions.

## Introduction

Human apoptosis, or programmed cell death, is mostly regulated by the B-cell lymphoma 2 (BCL-2) protein family.^1,2^ Many cancer types have overexpressed BCL-2 and its anti-apoptotic relatives such as BCL-XL, which makes tumor cells resistant to chemotherapy therapies.^1,3^ Consequently, BCL-2 has become a desirable target for designing and identifying innovative anti-cancer drugs.^1,2^ The BCL-2 protein family is essential for the prevention of cancer because it plays a critical role in the regulation of apoptosis.^4^ Many cancer forms are characterized by aberrant overexpression of BCL-2 family members, which block apoptosis, or by a reduction in pro-apoptotic proteins. Because of their critical function in regulating apoptosis, the pro-survival and pro-apoptotic BCL-2 family proteins are appealing targets for developing cancer treatment drugs.^5^ For efficient binding and inhibition of BCL-2, the ligand conformation within the protein binding site is critical. As prospective cancer therapies, small compounds that can bind to BCL-2 and cause conformational changes that impair the protein’s anti-apoptotic function are highly desirable. For structure-based drug design that targets BCL-2, precise ligand conformation prediction is crucial.^5,6^ The development of BCL-2 inhibitors and their discovery have advanced significantly in recent years.^2,7,8^ Targeting BCL-2 with BH3 mimetic small molecules has been verified with the FDA’s approval of Venetoclax, a selective BCL-2 inhibitor, in 2016 for the treatment of acute myeloid leukemia (AML) and chronic lymphocytic leukemia (CLL).^1,9^ Nevertheless, more effective and selective BCL-2 inhibitors with better therapeutic qualities and lower toxicity are still required.^1,3^ Discovering novel purposes for already-approved medications is known as drug repurposing, and it has shown promise in hastening the creation of cutting-edge cancer treatments.^2,7^ Researchers can find compounds with potential anti-cancer activity that have already undergone human safety testing by searching libraries of FDA-approved medications.^3^ Here, we report an *in silico* investigation employing conformations from the cutting-edge protein structure prediction tool NeuralPlexer^10^ to examine FDA-approved drugs that target the BCL-2 protein.^11^ A deep generative model called NeuralPlexer^10^ can simultaneously predict the three-dimensional (3D) structures of protein-ligand complexes directly from inputs such as protein sequences and ligand molecular graphs. In order to capture the intricate structural cooperativity between the protein and ligand, the model uses a hierarchical, multi-scale design to first predict residue-level contact mappings and then gradually refine the all-atom 3D coordinates.^12,13^ NeuralPlexer^10^ samples the ligand atoms’ 3D coordinates iteratively using a diffusion process, which enables the model to produce a variety of realistic and varied ligand conformations that meet important biophysical restrictions. NeuralPlexer’s diffusion process is engineered to include crucial biophysical priors, including bond lengths, angles, and steric conflicts, to facilitate the generation of physiologically plausible ligand conformations.^13^

In biological systems, protein-ligand interaction is ubiquitous and essential to comprehending cellular functions and creating efficient treatments. It has long been a major difficulty in computational biology and drug development to accurately anticipate the 3D structures of these protein-ligand complexes as well as the conformational changes brought about by ligand binding. This task has proven difficult for traditional computational techniques like physics-based methods such as molecular docking and molecular dynamics (MD) simulations, which frequently fail to reflect the initial intricate structural cooperativity between proteins and ligands. NeuralPlexer^10^ is a deep learning model that combines multiple state-of-the-art methods from generative AI, structural biology, and molecular modeling. The model uses a multi-scale, hierarchical design to predict residue-level contact maps first, followed by a step-by-step refinement of the all-atom 3D coordinates. Thus, NeuralPlexer^10^ can capture the intricate structural cooperativity that exists between the ligand and protein, which is a significant drawback of conventional docking methods that frequently regard the protein as rigid. The model can iteratively generate a variety of physically realistic ligand conformations that meet crucial biophysical constraints including bond lengths, angles, and steric clashes thanks to this diffusion process. Also, it can generate geometrically and energetically advantageous ligand conformations within the protein binding site by generating these biophysical priors directly into the generative process. The performance of this generative model has been thoroughly verified on numerous benchmarks, such as flexible binding site structure recovery and protein-ligand blind docking.^13^ The model routinely beats cutting-edge methods, including AlphaFold2, in terms of the precision of the global protein structure and the capacity to anticipate conformational changes brought on by ligands.^13^

To the best of our knowledge, this study is the first in the literature to utilize an FDA-approved drug library containing approximately 3,100 compounds to identify their binding sites within BCL-2 using NeuralPlexer. This process yielded around 1,300 successful binding poses, representing individual 3D conformations of BCL-2 for each bound ligand. Subsequently, we calculated the interaction energies (i.e., score-in-place) for these protein-ligand poses. Based on the interaction energy scores, further analyses were conducted, including binary-QSAR models to predict therapeutic activity (e.g., anticancer properties), as well as physics-based simulations and neural relational inference (NRI) analysis. Our results represent the first application of NeuralPlexer in drug repurposing, highlighting its effectiveness in accelerating the development of novel cancer therapeutics through *in silico* drug discovery techniques. Additionally, our findings provide valuable insights into the potential repurposing of existing drugs for the treatment of BCL-2-driven malignancies.

## Result and Discussion

In this study, we performed an *in silico* analysis to identify FDA-approved medications that show promise in targeting the BCL-2 protein, an important apoptosis regulator and a desirable target for cancer treatment. Utilizing the cutting-edge protein structure prediction model NeuralPlexer, we produced a library of 3,094 FDA-approved compounds as well as precise and sensitive conformations of the BCL-2 protein. Initially, we obtained the 3,094 FDA-approved compounds in .*SDF* format from the Drug Bank database. NeuralPlexer derived 8 conformations for every protein-ligand interaction using the whole sequence of the BCL-2 protein and then ranked them according to confidence scores. Therefore, we were able to concentrate on the most promising binding positions for additional examination. The protein-ligand complexes obtained for each BCL-2/ligand complex were scored by Glide SP (Standard Precision) with the “refine only” option which optimizes the ligand conformation without requiring any sampling. We were able to effectively assess the FDA-approved drugs’ binding to the BCL-2 protein binding site thanks to this method.

1,294 of the 3,094 drugs approved by the FDA were able to provide an acceptable output conformation during the refinement. We proceeded to further filter these compounds, choosing those that showed a good agreement between the two approaches (a root-mean-square deviation, or RMSD) value of less than 2 Å between the conformations generated by NeuralPlexer and refined conformation by Glide. Distribution of all 1,294 molecules’ docking scores and the proportion of compounds with RMSD values over and under 2 Å are shown in Figure 1A as a Violon plot.

**Figure 1.**
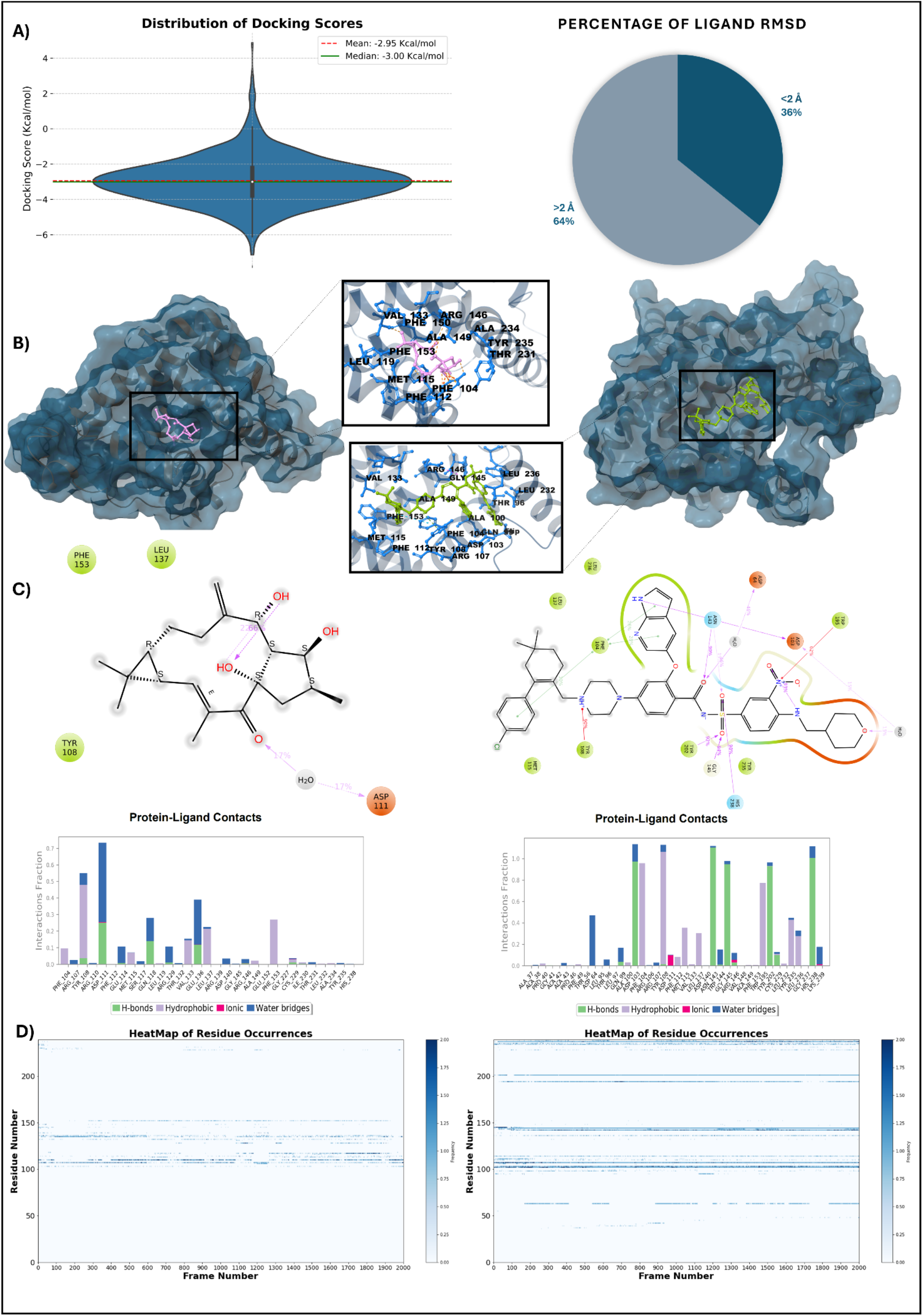
A) The Violon plot of the distribution of all 1294 molecules with the docking score and the percentage of molecules with RMSD over and under 2 Å; B) 3D structure of the BCL-2 protein in complex with Venetoclax in the bottom and Lathyrol in the top, along with their interacting residues around 3 Å; C) the interaction types of Venetoclax and Lathyrol in the bottom and top, respectively, during the 200 ns MD simulations; D) the heat map of the BCL-2 residues interacting with Venetoclax in the top and Lathyrol in the bottom.

After arranging the 1,294 docked compounds based on their Glide scores, the top-20 molecules were selected for further investigation. The highest-scoring (i.e., low binding energy) compounds were then subjected to a binary QSAR (Quantitative Structure-Activity Relationships) model using MetaCore (https://portal.genego.com/) to predict their anticancer activity. Out of the top-20, compounds with predicted anticancer activity scores greater than 0.5 were determined to be in the top-10, and the high Glide score was achieved for those compounds. The Desmond was then used to perform all-atom MD simulations (200 ns) on these 10 compounds, and the Molecular Mechanics/Generalized Born Surface Area (MM/GBSA) approach was used to determine the average binding free energies of the compounds. We also performed MM/GBSA computations and MD simulations for the FDA-approved BCL-2 inhibitor Venetoclax as a point of comparison. Venetoclax was shown to have an average MM/GBSA binding free energy of -148.8 kcal/mol, which is better than the top candidate compounds reported in our investigation. Since Venetoclax is a relatively large compound (i.e., molecular weight of 868 g/mol), in order to minimize the bias predicted biological activity, we normalized the GBSA scores with per nonhydrogen atom scores. Because it measures the binding energy per non-hydrogen atom by accounting for both the molecule’s size and binding affinity, ligand efficiency is a useful parameter in the drug discovery process. Improved solubility, permeability, and metabolic stability are among the advantageous drug-like characteristics that compounds with high ligand efficiency are typically more likely to possess. Five compounds outperformed Venetoclax in terms of ligand efficiency, which accounts for the number of non-hydrogen atoms in the molecule. These compounds were Lathyrol, Etofylline, Chlorzoxazone, Fadrozole, and Doxifluridine. Specifically, Lathyrol, Etofylline, Chlorzoxazone, Fadrozole, and Doxifluridine had average MM/GBSA scores of -43.56, -61.47, -35.75, -51.87, and -47.23 kcal/mol, respectively, and their corresponding ligand efficiency scores were -3.96, -3.84, -3.25, -3.05 and -2.78 kcal/mol. Due to their higher ligand efficiency scores, these substances may be able to attach to the BCL-2 protein binding site more successfully than Venetoclax which has ligand efficiency score of -2.44 kcal/mol, potentially resulting in increased potency and reduced off-target effects. Table 1 displays all the scores for the 10 FDA-approved drugs along with Venetoclax as the reference molecule.

**Table 1.**
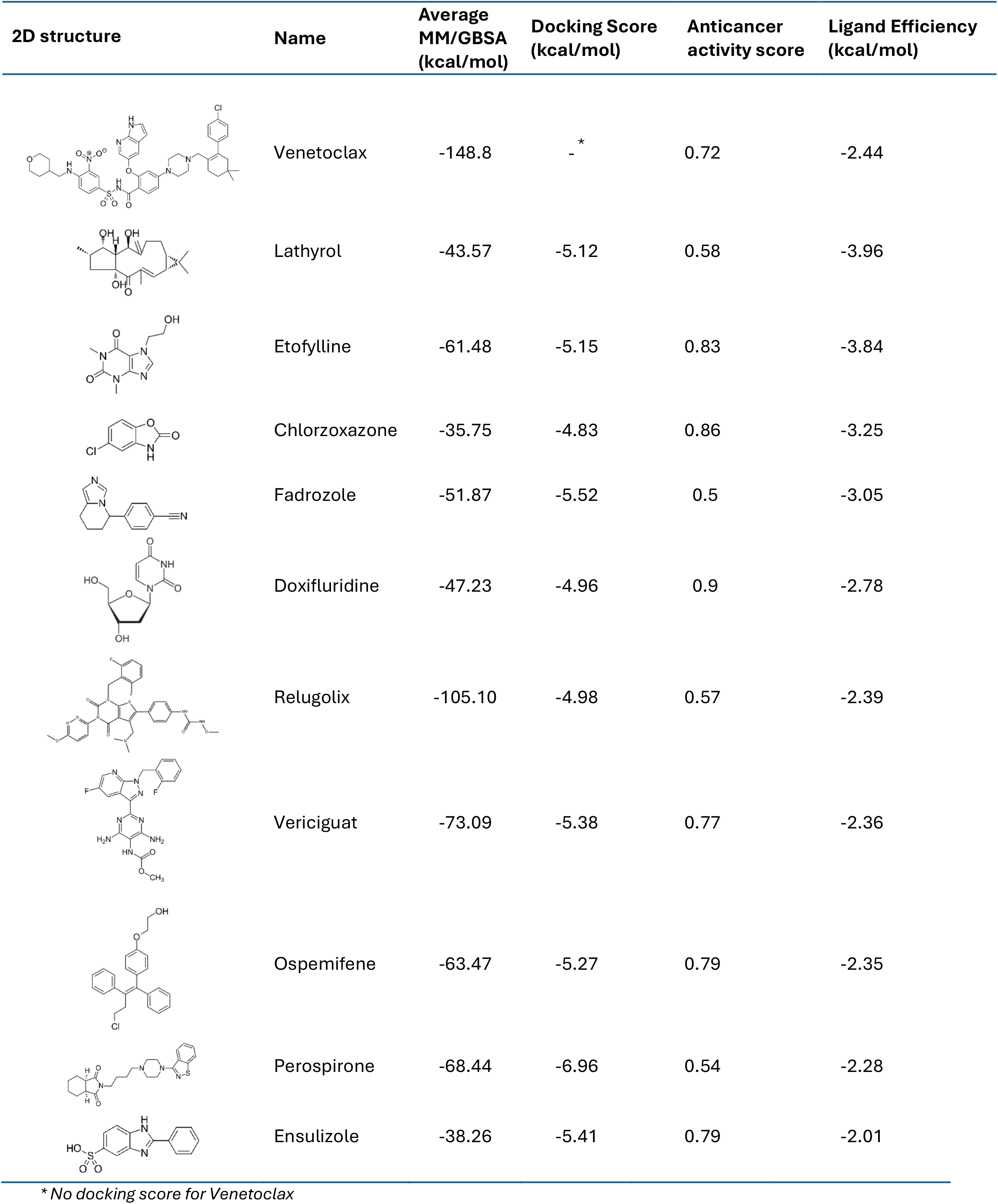
2D structures of 10 FDA-approved compounds with Venetoclax as the reference molecule, along with their MM/GBSA scores, docking scores, anticancer activity scores, and ligand efficiency statistics.

| 2D structure | Name | Average MM/GBSA (kcal/mol) | Docking Score (kcal/mol) | Anticancer activity score | Ligand Efficiency (kcal/mol) |
| --- | --- | --- | --- | --- | --- |
|  | Venetoclax | -148.8 | - * | 0.72 | -2.44 |
|  | Lathyrol | -43.57 | -5.12 | 0.58 | -3.96 |
|  | Etofylline | -61.48 | -5.15 | 0.83 | -3.84 |
|  | Chlorzoxazone | -35.75 | -4.83 | 0.86 | -3.25 |
|  | Fadrozole | -51.87 | -5.52 | 0.5 | -3.05 |
|  | Doxifluridine | -47.23 | -4.96 | 0.9 | -2.78 |
|  | Relugolix | -105.10 | -4.98 | 0.57 | -2.39 |
|  | Vericiguat | -73.09 | -5.38 | 0.77 | -2.36 |
|  | Ospemifene | -63.47 | -5.27 | 0.79 | -2.35 |
|  | Perospirone | -68.44 | -6.96 | 0.54 | -2.28 |
|  | Ensulizole | -38.26 | -5.41 | 0.79 | -2.01 |

\* No docking score for Venetoclax

The NRI model^14,15^ utilizes a variational graph autoencoder architecture to recreate the trajectories of MD simulations by learning latent contact maps between protein residues and ligand heavy atoms. While previous NRI models focused on understanding protein dynamics in apo and holo forms by tracking the carbon alpha atoms of amino acids, we have extended the NRI model to track both carbon alpha atoms and ligand heavy atoms. This enhancement allows us to quantify ligand-protein interactions, protein stability, and interactions between ligand heavy atoms. By training the NRI model to mimic MD trajectories, we obtain a latent interaction matrix *Z*_*ij*_ (0 ≤ *Z*_*ij*_ ≤ 1) that quantifies the contribution of atom *j* to the dynamics of atom *i*. The protein-ligand interaction energy is evaluated using the matrix *Z*_*ij*_, with details provided in the Methodology section. We trained 10 NRI models for 10 candidate compounds and Venetoclax, using the same hyperparameters for the variational graph autoencoder models. Figure 2A illustrates the correlation between protein-ligand interaction energy and ligand efficiency scores for these 10 compounds, yielding a Pearson correlation coefficient of r=*−*0.59. This negative correlation suggests that compounds with higher energy scores (indicating higher stabilization of protein dynamics^25^) tend to have lower ligand efficiency scores. Our NRI analysis thus supports the physics-based ligand efficiency scores, with both approaches affirming the potency of the compound Lathyrol. Figure 2B represents the interaction matrices *Z*_*ij*_ for Lathyrol and the reference molecule Venetoclax. The protein-ligand interaction maps indicate that the heavy atoms of Lathyrol significantly affect residue dynamics, resulting in higher normalized protein-ligand interaction energy compared to Venetoclax. Similarly, Figure 2B shows that the mean energy contribution of Lathyrol’s heavy atoms to each protein residue is generally higher (except for residue 81) than that of Venetoclax’s heavy atoms.

**Figure 2.**
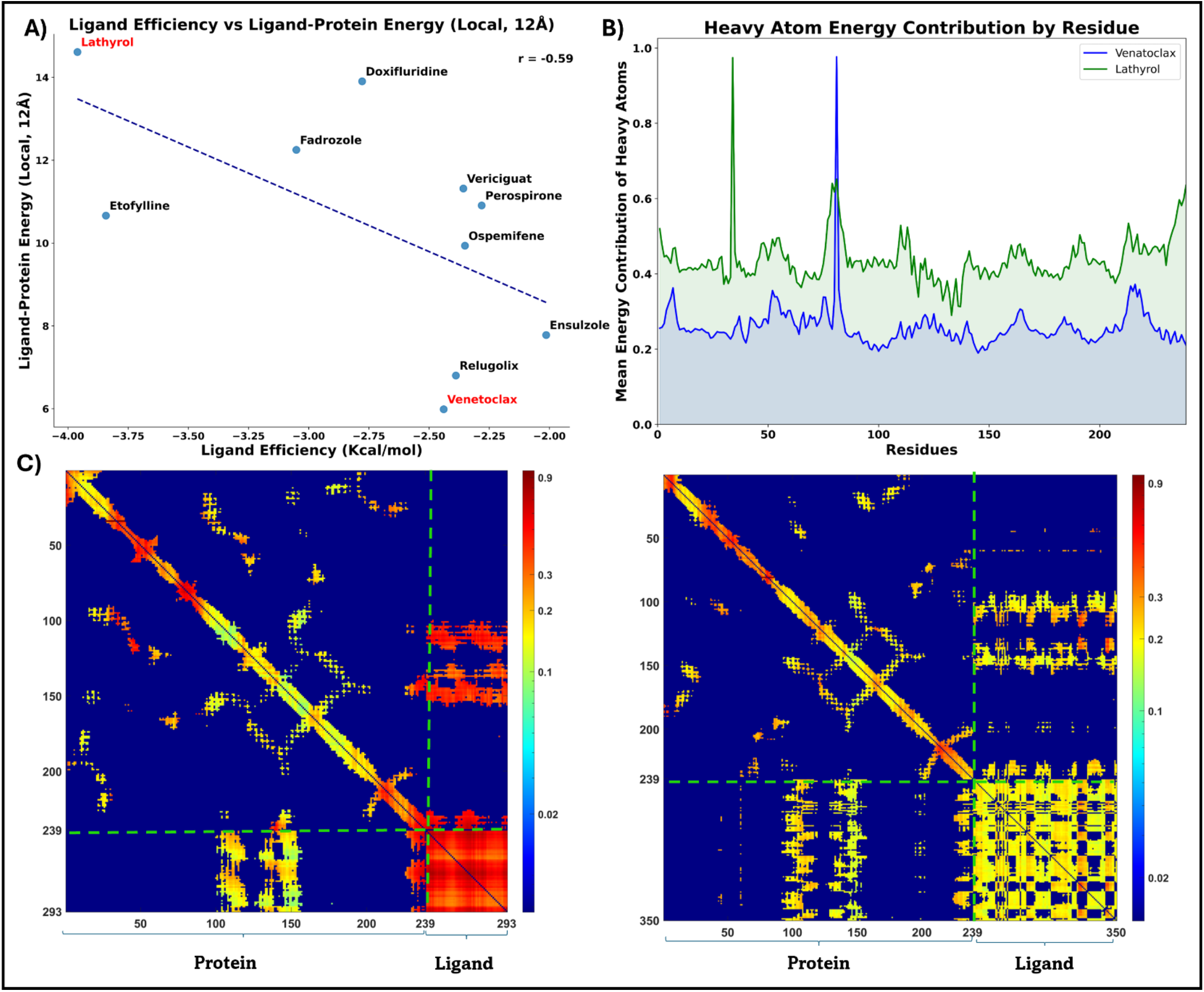
The results of the Neural Relational Inference (NRI) model are shown as follows: A) a scatter plot of the contributions of all ligands with a 12 Å radius around the BCL-2 protein residues and the compounds’ ligand efficiency (Chlorzoxazone removed from the plot as an outlier); B) the mean energy contribution of the heavy atoms of Lathyrol and Venetoclax with BCL-2 protein residues; C) a contact map of BCL-2 with Lathyrol on the left, and a contact map of BCL-2 with Venetoclax on the right.

## Discussion

The outcomes of this *in silico* investigation demonstrate the value of combining cutting-edge protein structure prediction software (i.e., NeuralPlexer) with conventional computational drug discovery techniques. Through the utilization of NeuralPlexer’s precision in predicting protein-ligand complex structures, we effectively screened an extensive collection of FDA-approved drugs and detected potential candidates for BCL-2 inhibitors. Based on their favorable binding free energies and high ligand efficiency ratings, Lathyrol, Etofylline, Chlorzoxazone, Fadrozole, and Doxifluridine were identified as the top-performing compounds. This suggests that these molecules should be investigated further as potential cancer therapies targeting BCL-2. These compounds’ enhanced ligand efficiency over the FDA-approved Venetoclax underscores the possibility of drug repurposing to produce more effective and specific BCL-2 inhibitors.

The transformational impact of modern protein structure prediction tools is demonstrated by the successful integration of NeuralPlexer into our computational drug development approach. NeuralPlexer allowed us to construct high-quality conformations that were essential for the subsequent physics-based binding free energy calculations by precisely capturing the complex structural cooperativity between proteins and ligands.

The oral anticancer drug Doxifluridine had an average MM/GBSA score of -47.23 kcal/mol and a ligand efficiency score of -2.78 kcal/mol. It also has antiangiogenic properties. Even if it’s not one of the best options, Doxifluridine is a worthwhile substance to look into further in the context of BCL-2-driven tumors due to its capacity to suppress angiogenesis and cause apoptosis in cancer cells. After Doxifluridine, the ligand efficiency of Fadrozole, an aromatase inhibitor, is superior at -3.05 kcal/mol. It is approved to treat breast cancer and has an average MM/GBSA score of -51.87 kcal/mol. This implies that Fabrozole might be able to maintain a relatively modest molecular size while exhibiting a high affinity for binding to the BCL-2 protein binding site. Following Fadrozole, Chlorzoxazone has a slightly better ligand efficiency score of -3.25 kcal/mol. The possibility of the muscle relaxant chlorzoxazone as a BCL-2 inhibitor has not been previously explored. With an even higher ligand efficiency score of -3.84, etofylline, a phosphodiesterase inhibitor used to treat cerebrovascular diseases, shows promise as a strong and selective BCL-2 inhibitor. Among the best choices, Lathyrol, a naturally occurring substance present in several legumes, has promising ligand efficiency score (−3.96 kcal/mol). Even though Lathyrol is not a recognized medication, its advantageous binding properties and room for improvement make it a promising lead molecule for future research.

It is crucial to remember that this study’s findings are based on *in silico* predictions and need to be confirmed through experiments. Nonetheless, a strong framework for identifying potential therapeutic candidates is provided by the combination of NeuralPlexer’s precise prediction of protein-ligand complex structure and the ensuing physics-based binding free energy calculations. The discovery that Lathyrol, Etofylline, Chlorzoxazone, Fadrozole, and Doxifluridine are the best-performing compounds with the highest ligand efficiency scores indicates that these compounds should be studied further in *in vitro* studies.

In summary, this study highlights the need of merging state-of-the-art protein structure prediction instruments, such as NeuralPlexer, with traditional computational drug discovery methodologies, and it is found that certain compounds are found to have the ability to inhibit BCL-2 more effectively than Venetoclax, based on their ligand efficiency. To further support our physics-based observations, we extended the scope of the NRI model to measure protein-ligand interaction patterns. The novel NRI model can quantify residue-residue, ligand-residue, and ligand-ligand contacts, providing an energy/stability score that reflects how ligand and residue atoms influence the protein/ligand structure throughout the simulation. The compounds’ ligand efficiency and NRI protein-ligand interaction scores show a negative correlation (r = -0.59, Figure 2A). This indicates that the global NRI analysis (trained with all-atom trajectories) aligns with the local ligand efficiency scores. After examining the top 5 molecules that outperformed Venetoclax in terms of ligand efficiency, we discovered that two of them had the highest energy ratings of all the molecules. This indicates that these compounds increase the stability of the BCL-2 protein in comparison to Venetoclax. With a ligand-protein energy score of 104.006 and 97.17, respectively, Lathyrol and Fadrozole stabilize the protein dynamics much more than the others. These findings strongly imply that Lathyrol and Fadrozole are promising BCL-2 inhibitors that could perform better than Venetoclax. Lathyrol and Fadrozole exhibit high ligand efficiency and NRI stability ratings, suggesting that they can bind to the BCL-2 protein binding site with remarkable affinity while retaining a comparatively tiny molecular size. Better solubility, permeability, and metabolic stability are just a few of the drug-like qualities that might result from having this desirable trait. The results of this thorough *in silico* study demonstrate Lathyrol and Fadrozole’s promise as new BCL-2 inhibitors that call for more experimental validation and refinement. Based on their better performance than the FDA-approved Venetoclax, these compounds may be developed into more potent and selective cancer therapies that target the BCL-2 pathway. The NRI analysis of the compound Lathyrol has demonstrated its potential as a cancer-targeting inhibitor (Figures 2B and 2C).

The success of NeuralPlexer in accurately predicting BCL-2 co-crystallized structures was also investigated. We used the 6O0K PDB coded structure for comparison. The RMSD between the protein structures was measured as 2.47 Å, and the RMSD of Venetoclax conformers within the binding pocket was measured as 0.35 Å. (Figure 3) The results confirm that NeuralPlexer accurately predicts the ligand-bound conformers of protein-ligand complex structures. This capability may allow researchers to bypass traditional docking processes, which can sometimes yield misleading results, and instead start simulations and analyses with correct ligand-bound forms.

**Figure 3.**
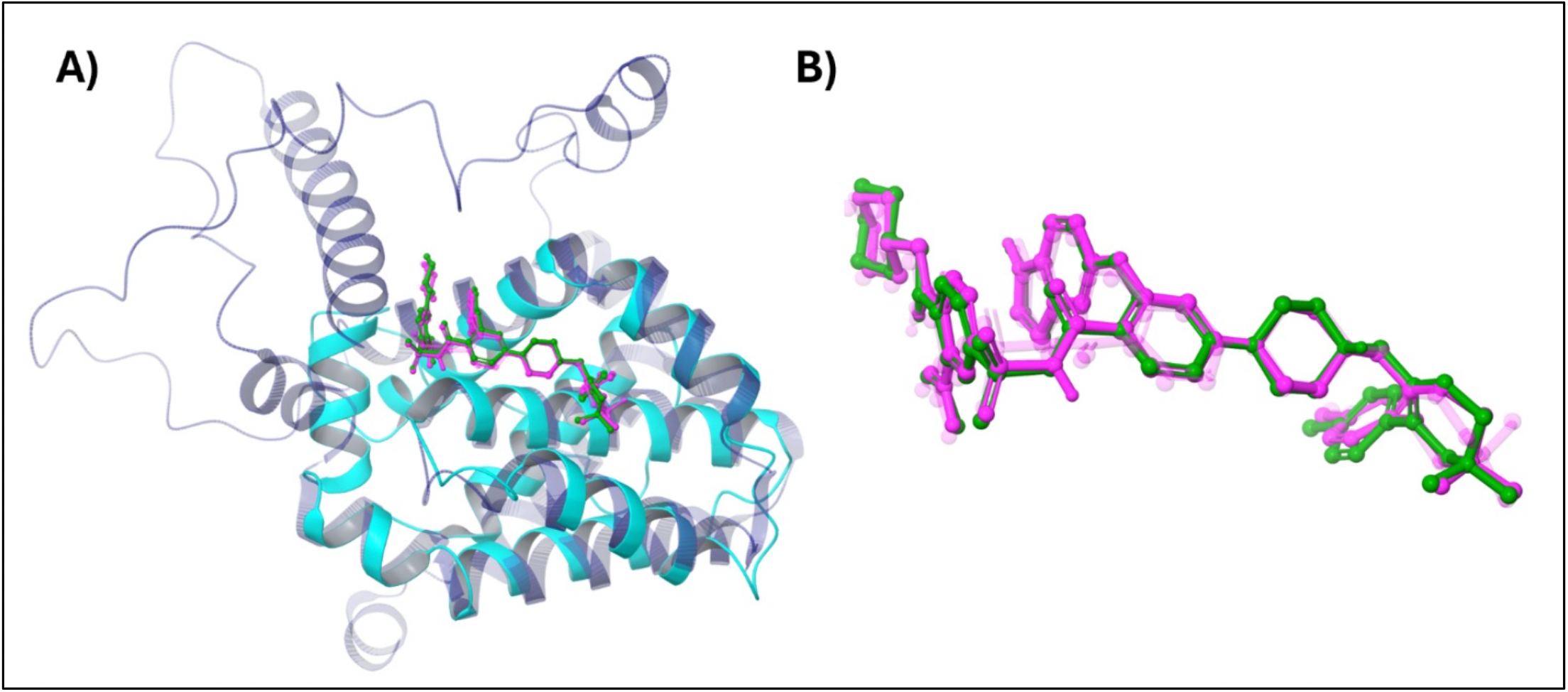
Comparison of co-crystallized BCL-2 structure of Venetoclax (PDB: 6O0K) with NeuralPlexer predicted 3D structure. X-Ray structure of BCL-2 is colored with cyan while NeuralPlexer predicted structure represented with ice-blue. Venetoclax is colored as pink (X-ray conformer) and green (NeuralPlexer prediction)

## Methods

### BCL2-Ligand Complex Conformations

In this study, we generated high-quality conformations of the BCL-2 protein and the 3,094 FDA-approved drugs from the DrugBank database using the cutting-edge protein structure prediction tool, NeuralPlexer^10^. Using protein sequences and ligand molecules as input, the deep generative model NeuralPlexer can accurately predict the 3D structures of protein-ligand complexes. The following are the main ways that NeuralPlexer^10^ creates the conformations: NeuralPlexer^10^ samples the 3D coordinates of the ligand atoms iteratively by means of a diffusion process. This enables the model to provide realistic and varied ligand conformations that meet important biophysical requirements. Also, the model uses a hierarchical multi-scale architecture to improve the all-atom 3D coordinates step-by-step after predicting residue-level contact maps. NeuralPlexer^10^ is able to capture the intricate structural cooperativity between the ligand and protein as a result. NeuralPlexer’s diffusion mechanism is engineered to take into account crucial biophysical limitations such link lengths, angles, and steric conflicts. This aids in the generation of physiologically plausible ligand conformations by the model. NeuralPlexer^10^ can capture the conformational changes brought about by ligand binding since it is trained to sample the protein in both its ligand-bound and ligand-free states. An ensemble of predicted ligand conformations is produced by NeuralPlexer^10^; these can be sorted and chosen according to confidence scores or other standards to determine the most promising binding poses. The following particular settings that are applied when using NeuralPlexer: Produce 10 distinct forms for every protein-ligand combination, divide the input into 4 segments to maximize memory utilization, 40 diffusion stages are needed to produce the final conformations, utilize a GPU to run the model for quicker computation, apply the sampling strategy of Langevin Simulated Annealing, find and manage covalent connections between the ligand and the protein, and sort the produced conformations according to the scores for confidence. NeuralPlexer^10^ produced high-quality conformations, which served as the foundation for further computational investigation. This included calculations of binding free energy, molecular docking, and MD simulations. The success of this investigation was largely dependent on NeuralPlexer’s capacity to precisely capture the intricate structural cooperativity between the BCL-2 protein and small molecule ligands.

### Binding Score Predictions

The protein-ligand complex conformations produced by NeuralPlexer^10^ offer insightful structural information, but predicted binding affinities must be determined in order to properly appraise these complexes’ potential as therapeutic candidates. In order to do this, we created a Python script that makes use of the Schrödinger’s Maestro software suite’s command-line interface (which is accessible through our GitHub repository at www.github.com/DurdagiLab

The script uses Maestro’s built-in features to merge and prepare the protein-ligand complex structures produced by NeuralPlexer ^10^. It then generates a grid box centered on the centroid of the ligand, defining the binding site for the docking calculations. Subsequently uses Glide, to forecast each complex’s Standard Precision (SP) docking scores. Following that, using Glide’s “refine only” option, which, without requiring comprehensive sampling, quickly optimizes the ligand conformation from the pose provided by NeuralPlexer^10^ and reports the optimized Glide SP docking scores for each protein-ligand complex conformation generated by NeuralPlexer.

Through the integration of Glide’s dependable docking and scoring capabilities with NeuralPlexer’s precise complex structure prediction, we can effectively assess the predicted binding affinities of several possible hit candidates. Glide’s “refine only” option strikes a compromise between computational efficiency and binding score prediction accuracy by permitting ligand conformational modifications without compromising the overall binding pose predicted by NeuralPlexer ^10^.

By combining the benefits of Maestro’s and NeuralPlexer’s computational tools, this integrated strategy helps researchers quickly screen and rank drug candidates according to their predicted binding affinities. The Python script accelerates the drug development process by streamlining the workflow and enabling high-throughput analysis of huge chemical libraries.

### Binary QSAR Anticancer Activity Score

The molecules were organized into separate .*sdf* files according to their docking scores once their SP docking scores were determined and, when using the “refine only” option, their conformation change was not greater than 2 Å. After that, these files were uploaded to the MetaCore™ ^16^ platform, so that the “cancer therapeutic activity prediction QSAR” feature could be used for screening. This characteristic makes it possible to predict a compound’s likelihood of having anticancer properties. The Cancer-QSAR model predicts the compounds’ therapeutic activity values (TAV) by comparing the input compounds to those with established high anticancer activity during the screening process. TAV levels are standardized between 0 and 1, with values greater than 0.5 suggesting possible anticancer action. Using a training set of 886 compounds and a set of descriptors, the cancer-QSAR model in the MetaCore™ ^16^ was developed, yielding a sensitivity of 0.95, specificity of 0.92, accuracy of 0.93, and Matthew’s Correlation Coefficient (MCC) of 0.87. A test set of 167 chemicals was used to further validate the model, and results showed a sensitivity of 0.89, specificity of 0.83, accuracy of 0.86, and MCC value of 0.72. An initial screening result of 0.5 was used to calculate the cutoff threshold. All other compounds were filtered out and only the top 10 molecules with the highest docking score between those with a predicted cancer TAV of more than or equal to 0.5 were chosen for additional analysis.

### Structure Refinement

ModRefiner^17^, a high-resolution protein structure refinement server created by the Yang Zhang Lab, was used to further refine the molecules that met the binary QSAR score criterion of 0.5. Starting from C-alpha traces, main-chain models, or full-atomic models, the ModRefiner ^17^ program can refine protein structures down to the atomic level. A combination of knowledge- and physics-based force fields steers the conformational search while the sidechain and backbone atoms remain fully flexible during the refining process. In addition, ModRefiner^17^ provides the ability to steer the refinement simulations using a reference structure, bringing the basic starting models closer to their native state in terms of side-chain placement, backbone topology, and hydrogen bonding. Furthermore, ModRefiner^17^ produces notable enhancements in the physical attributes of nearby structures. *Ab initio* full-atomic relaxation, in which the improved model is unrestricted by the original or reference models, is supported by the stand-alone application. This step was required since there could be some proline structural mistakes in the protein structures that were obtained from NeuralPlexer ^10^.

### Structure Preparation

To prepare the protein, the Protein Preparation Tool ^18^ was used. This included reassembling disulfide bonds, adding hydrogen atoms, forming zero-order bonds with metals, and allocating bond orders. Water molecules that were more than 5 Å from hetero groups were removed. With the help of PROPKA, the target protein’s residues’ protonation states at physiological pH were assigned, and the OPLS3e force field ^19^ was used to reduce the side chain atoms.

### MD simulations

All-atom MD simulations are the next step, following the preparation of the top 10 compounds and Venetoclax as the reference molecule. The protein-ligand complex was positioned in a solvation box with the TIP3P water ^19^model in the simulations, which were run using the Desmond^20^. A buffer zone measuring 10 Å in an orthorhombic box encircled the compound. The NPT ensemble was used with a constant pressure of 1.01325 bar and a temperature of 310 K in order to attain thermodynamic equilibrium. Na^+^ ions were used to neutralize the system at first, and 0.15 M NaCl was supplied to keep the pH of the system at 7.4. Using the Nose-Hoover thermostat^21^ and Martyna-Tobias-Klein barostat^22^, each compound was simulated separately. The RESPA integrator was used with various time steps to manage bonded, non-bonded, and distant nonbonded interactions. Long-range electrostatic interactions were estimated using the particle mesh Ewald method^23^ and periodic boundary conditions (PBC), whereas short-range electrostatics and van der Waals interactions were calculated with a cutoff value of 9 Å. The system’s potential energy was ascertained using the OPLS3e force field. Desmond was used to reduce and relax the simulation box before the simulations. For further analysis, a set of 2000 frames were extracted from every 200 ns in these simulations.

### MM/GBSA Binding Free Energy Calculations

The MM/GBSA approach was utilized with the Prime^24^ module of Maestro to calculate the average binding energies. 2000 protein-ligand complexes were collected from the trajectories of each MD simulation running for 200 ns. The system’s dielectric constant could vary since the VSGB 2.0 ^25^ implicit solvation model was applied. The system’s external dielectric constant, 80, was viewed as constant, but the system’s internal dielectric constant may vary between 1.0 and 4.0 under the OPLS3e force field.^19^ The average values and standard deviations were computed for each compound after the MM/GBSA for each complex was determined.

### NRI Model Analysis

It is crucial to comprehend the intricate relationships found in nature, particularly as they relate to physical dynamical systems. Complex interacting atoms are produced by MD simulations, and conventional examination of pairwise relationships between residues requires the assumption of linear correlation. The NRI model has an unsupervised variational graph-based autoencoder architecture^14^ and have been proposed for the investigation of complex interactions in dynamical systems, particularly in protein dynamics.^15^ By comprehending the nonlinear linkages as an interaction graph or latent representation *Z*_*ij*_, which represents the strength of the interaction between residues *i* and *j*, the NRI model seeks to recover dynamic trajectories. To evaluate protein-ligand energy scores, we use the formula:

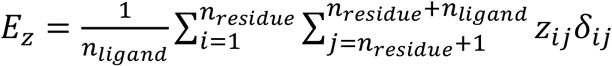

where *n*_*residue*_ and *n*_*ligand*_ represent protein residue number and ligand heavy atom number, *Z*_*ij*_ is the learned interaction matrix and *δ*_*ij*_ = 1 for 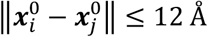 and *δ*_*ij*_ = 0 for 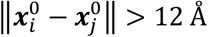. We followed the literature to select 12 Å as the threshold distance while evaluating the energy score *E*_*Z*_.^15^ Higher energy scores meaning that the ligand more stabilizes the protein function. In other words, *E*_*Z*_ measures how much ligand’s heavy atoms affect the residue dynamics in average.

In this study, we further developed the NRI model to regenerate not only residue trajectories but also ligand trajectories by learning complex residue-residue, residue-ligand and ligand-ligand interactions. We examined the 2000 frame MD trajectories of 11 compounds in total that were complex with the BCL-2 protein. The literature contains all the mathematical details of the NRI model as well as its source codes.^15^ We have redesigned source codes^15^ of the NRI model to work with trajectories of protein-ligand complexes. To speed up and automate the process-the revised NRI for protein-ligand complexes codes have been uploaded to our Github page (www.github.com/DurdagiLab). It gathers the files, initiates NRI model, analyzes the interaction maps, and saves them as plots and CSV files.

